# Does force depression resulting from shortening against series elasticity contribute to the activation dependence of optimum length?

**DOI:** 10.1101/2024.06.06.597766

**Authors:** Dean L Mayfield, Natalie C Holt

**Affiliations:** Department of Evolution, Ecology, and Organismal Biology, University of California, Riverside, Riverside, California, USA

**Keywords:** History dependence, series compliance, force-length relationship, maximum force, myofilament overlap, stretch-induced muscle damage

## Abstract

The optimum length for force generation (*L*_0_) increases as activation is reduced, challenging classic theories of muscle contraction. Although the activation dependence of *L*_0_ is seemingly consistent with length-dependent Ca^2+^ sensitivity, this mechanism can’t explain the apparent force dependence of *L*_0_, or the effect of series compliance on activation-related shifts in *L*_0_. We have tested a theory proposing that the activation dependence of *L*_0_ relates to force depression resulting from shortening against series elasticity. This theory predicts that significant series compliance would cause tetanic *L*_0_ to be shorter than the length corresponding to optimal filament overlap, thereby increasing the activation dependence of *L*_0_. We tested this prediction by determining *L*_0_ and maximum tetanic force (*P*_0_) with (*L*_0_spring_, *P*_0_spring_) and without added compliance in bullfrog semitendinosus muscles. The activation dependence of *L*_0_ was characterised with the addition of twitch and doublet contractions. Springs attached to muscles gave added fixed-end compliances of 11-39%, and this added compliance induced force depression for tetanic fixed-end contractions (*P*_0_spring_/*P*_0_ < 1). We found strong, negative correlations between spring compliance and both *P*_0_spring_ (*r*^2^ = 0.89-91) and *L*_0_spring_ (*r*^2^ = 0.60-63; *P* < 0.001), while the activation dependence of *L*_0_ was positively correlated to added compliance (*r*^2^ = 0.45, *P* = 0.011). However, since the compliance-mediated reduction in *L*_0_ was modest relative to the activation-related shift reported for the bullfrog plantaris muscle, additional factors must be considered. Our demonstration of force depression under novel conditions adds support to the involvement of a stress-induced inhibition of cross-bridge binding.

## INTRODUCTION

The isometric force-length relationship of skeletal muscle (Gordon et al., 1966; Edman & Reggiani, 1987; Winters et al., 2011; Moo et al., 2020) is largely consistent with the sliding filament and cross-bridge theories of contraction, which give relatively simple structural and molecular mechanisms of force generation (Huxley & Hanson, 1954; Huxley & Niedergerke, 1954; Huxley, 1957). Together, these theories posit that muscle length change and force generation are due to the relative sliding of myosin (i.e., thick) and actin (i.e., thin) filaments in response to cyclical interactions of myosin-based cross-bridges with attachment sites on actin. The net contractile force resulting from myosin-actin interaction is dictated by the force generated per cross-bridge and the number of cross-bridges, which varies as a function of the overlap between thick and thin filaments. Myofilament overlap and, therefore, force, are maximal over a small range of intermediate sarcomere lengths. Whilst the predictions of these classic theories of muscle contraction generally hold true for steady-state force in maximally activated muscle, they are often violated for conditions of submaximal activation. Numerous studies have shown that the optimum length for force generation progressively increases when muscle activation is gradually reduced (Rack & Westbury, 1969; Roszek et al., 1994; Balnave & Allen, 1996; Brown et al., 1999; Morgan et al., 2000), sometimes by more than 30% (Holt & Azizi, 2014, 2016).

In this context, submaximal activation refers to incomplete activation of the thin filament through the binding of calcium ions to the regulatory protein, troponin C. When the free concentration of intracellular calcium is subsaturating (i.e., submaximal), actin-myosin interaction remains partially inhibited by the regulatory protein tropomyosin such that force generation is submaximal. The relationship between free Ca^2+^ concentration and thin filament activation, or force, is modulated by sarcomere length, possibly through a mechanism related to filament lattice spacing (Allen & Moss, 1987; Martyn & Gordon, 1988; Wang & Fuchs, 1995; Rockenfeller et al., 2022). With increasing sarcomere length, the sigmoidal relationship between free Ca^2+^ concentration and force shifts leftward, indicating elevated Ca^2+^ sensitivity at longer sarcomere lengths (Stephenson & Williams, 1982; Stienen et al., 1985; Martyn & Gordon, 1988). This length dependence in the Ca^2+^ sensitivity of thin filament activation can be easily reconciled with the activation dependence of optimum length. In skinned muscle fibres, where the concentration of free Ca^2+^ is directly controlled, elevated Ca^2+^ sensitivity at long lengths more than compensates for the loss of force-generating potential from reduced filament overlap; optimum sarcomere length increases with decreasing activation (Moss et al., 1983; Stephenson & Wendt, 1984; Stienen et al., 1985; de Beer et al., 1988).

In an intact fibre or muscle, Ca^2+^ release is triggered by membrane depolarisation such that Ca^2+^ availability and thin filament activation are modulated by the frequency of excitation (Blinks et al., 1978; Westerblad et al., 1993; Balnave & Allen, 1996). In agreement with observations on skinned fibres, lowering activation by decreasing stimulation frequency causes optimum length to increase (Rack & Westbury, 1969; Balnave & Allen, 1996; Zuurbier et al., 1998; Brown et al., 1999; Morgan et al., 2000). Additional support for the involvement of length-dependent Ca^2+^ sensitivity comes from studies demonstrating that the activation related-shift of optimum length is less pronounced following post-activation potentiation (Brown & Loeb, 1998), which increases Ca^2+^ sensitivity (Sweeney et al., 1993), and arises for tetanic stimulation following treatment with sodium dantrolene (Wendt & Barclay, 1980), which reduces Ca^2+^ release from the sarcoplasmic reticulum (Desmedt & Hainaut, 1977).

Activation and force are inextricably linked, and until more recently, these two factors had not separated. At the level of the whole muscle, force also varies according to motor unit recruitment. An argument centred on force transmission and internal work was formulated since optimum length was found to be independent of thin filament activation (i.e., Ca^2+^ concentration) but dependent on muscle force (Holt & Azizi, 2014). Muscle force was varied by lowering muscle recruitment (i.e., stimulation intensity) rather than activation at the level of the fibre (i.e., stimulation frequency). Using tetanic stimulation over a range of submaximal stimulation voltages, optimum length was shown to increase with decreasing muscle force. This result was attributed to the multiple sources of compliance within the force transmission pathway that necessitate internal work be performed by the contractile apparatus for force to be registered at the attachment points of the muscle. For low-force contractions, this internal work requirement is disproportionately greater. Fewer cross-bridges must do the same amount of work on the elastic force-transmitting structures. Increasing muscle length, however, can reduce internal work by pre-tensioning these structures. As such, force generation is favoured at relatively long muscle lengths for low-force contractions despite less-than-optimal filament overlap. Although related, this mechanism is not to be confused with the nonlinear load-extension relationship of tendon and aponeurosis (e.g., Bennett et al., 1986; Scott and Loeb, 1995) that can exploited for brief contractions by increasing muscle length and passive tension such that series elastic stiffness is elevated at contraction onset. Greater series elastic stiffness will increase the rate of force development (e.g., Hill, 1951) but can’t directly improve steady-state force.

If the requirement for internal work in force transmission pathways is the principal cause of activation-related shifts in optimum length, increasing muscle length beyond optimum filament overlap should offer the greatest advantage when muscle recruitment is reduced to a single motor unit, but this is not the case. Even though the average single motor unit force may be less than 1% of whole muscle tetanic force (Lewis et al., 1972; Bagust et al., 1973; Stephens et al., 1975), the optimum muscle length for a twitch or tetanus of a single motor unit can be similar or even shorter than the optima of the parent muscle (Lewis et al., 1972; Bagust et al., 1973; Stephens et al., 1975; Luff & Proske, 1979; Filippi & Troian, 1994). Differences in mean sarcomere length across motor units and with respect to the whole muscle could help account for this observation. However, since the force generated by a fraction of muscle (i.e., small group of motor units, ventral root filament or bundle) appears to be constrained only when the volume of muscle recruited is relatively small, and only by a small amount (Brown & Matthews, 1960; Emonet-Dénand et al., 1990; Perreault et al., 2003; Drzymała-Celichowska et al., 2010), the requirement to do work on force-transmitting structures seems unlikely to explain large shifts in optimum length.

An alternative interpretation of the force dependence of optimum length demonstrated by Holt and Azizi (2014) has been formulated on the basis that series elastic compliance permits active shortening during force development and active shortening induces force depression (Holt & Williams, 2018). This idea is supported by the observation that the activation dependence of optimum fibre length is considerably more pronounced for the bullfrog plantaris muscle than it is for a fibre bundle from this muscle (Holt & Williams, 2018). Because substantial series elastic compliance resides in aponeuroses and external tendons, active fibre shortening is seemingly unavoidable for an isometric or, rather, fixed-end contraction of an intact muscle (Griffiths, 1991; Scott & Loeb, 1995; Buchanan & Marsh, 2001; Ahn et al., 2003; MacIntosh & MacNaughton, 2005; Azizi & Roberts, 2010; Mendoza & Azizi, 2021). For the bullfrog plantaris muscle, active shortening against the stretch of series elasticity amounts to roughly 30% of fibre length (Azizi & Roberts, 2010; Sawicki et al., 2015).

Shortening-induced force depression refers to the deficit in steady-state isometric force that arises at a given length after a period of active shortening. Conventionally, shortening is imposed upon the muscle as force approaches a maximum amplitude or shortly afterwards. Although the mechanism underpinning force depression is not agreed upon (Rassier & Herzog, 2004; Nishikawa et al., 2012, 2018; Holt & Williams, 2018), it is clear that the magnitude of force depression after active shortening depends strongly on the amount of mechanical work performed during shortening (Granzier & Pollack, 1989; Herzog et al., 2000; Lee & Herzog, 2009). Because active shortening against series elasticity increases in proportion to force, and force increases in proportion to activation (and recruitment), the mechanical work performed during a fixed-end contraction will increase in proportion to activation. If we approximate the load-extension relationship of the series elastic element to be roughly linear, the amount of work performed during force development will be proportional to shortening squared such that force depression should vary relatively more than force as activation is modulated.

The work dependence of force depression alone is insufficient to facilitate a force dependence of optimum length. Critically, force depression exhibits a dependence on length (Edman et al., 1993; Meijer et al., 1998; Morgan et al., 2000). Force depression normalised to mechanical work is more pronounced at longer muscle lengths, increasing in an exponential fashion (Van Noten & Van Leemputte, 2011). This length dependence provides a means for force depression to alter the shape of the force-length relationship. This theory, which we will refer to as the *force depression hypothesis*, posits that the activation or force dependence of optimum length arises, at least in part, through the interaction of the work and length dependence of force depression and the length dependence of myofilament overlap (i.e., force generating potential). That is, as force increases in response to increases in activation (and recruitment), so does force depression, and because force depression is disproportionately greater at longer lengths, optimum length becomes increasingly shorter with increasing activation. This theory predicts that the length corresponding to maximum tetanic force is shorter than the length corresponding to optimal filament overlap. By contrast, mechanisms centred on the length dependence of Ca^2+^ sensitivity and internal work do not cause tetanic optimum length to deviate from optimum filament overlap. Instead, the activation dependence of optimum length arises because optimum length becomes progressively longer with respect to optimum filament overlap as activation is reduced.

The *force depression hypothesis* puts forward at least two testable predictions: 1) tetanic optimum length (*L*_0_) can be reduced though an increase in series elastic compliance, and 2) the activation or force dependence of *L*_0_ can be increased through an increase in series elastic compliance. If prediction 1 is confirmed, and the reduction is of sufficient magnitude to explain prior observations, it seems unlikely that the requirement for internal work in force transmission pathways is responsible for the apparent force dependence of *L*_0_ because the involvement of this mechanism requires that the deviation from optimal filament overlap occurs for low-force contractions and in the direction of longer lengths. We tested the aforementioned predictions by constructing twitch, doublet, and tetanic force-length relationships for isolated muscles with and without added in-series compliance, which was given by a linear extension spring. Spring stiffness was varied between muscles to give a range of effective compliances. Because this study serves as the first to directly characterise force depression during the development of tetanic force it may also offer new insights into the mechanism responsible.

## MATERIALS AND METHODS

### Animals

Adult bullfrogs (*n* = 17), *Lithobates catesbeianus*, were purchased from a herpetological vendor and ranged in body mass from 63 to 157 g. Animals were housed in large aquaria that provided both aquatic and terrestrial regions and were fed large crickets *ad libitum* twice weekly. All animal husbandry and experimental procedures were approved by the Institutional Animal Care and Use Committee of the University of California, Riverside (Experiment Protocol: 20220027).

### *In vitro* muscle preparation

Frogs were anaesthetised with vaporised isoflurane and euthanised by a double pithing protocol. Frogs were placed in a dissection tray containing chilled amphibian Ringer’s solution that lay on a bed of ice. The skin overlaying the right leg was removed and the dorsal head of the semitendinosus muscle was isolated using blunt dissection. The distal tendon was knotted with Kevlar and severed at the point where the tendon begins to fan out before inserting onto the tibiofibula. A small stainless-steel jump ring (diameter: 4 mm; mass: 29 mg) was secured to this distal tendon using Kevlar to allow the muscle to be connected to a spring. The proximal tendon was knotted with Kevlar at the insertion of the tendon onto the pelvis before a small segment of pelvis was cut free. Two small knots of 6-0 silk suture were placed towards each end of a superficial fascicle and served as markers for measuring fascicle segment length.

The muscle was placed in a horizonal Perspex muscle bath between two platinum plate electrodes connected to a universal isolated stimulator (UISO, Harvard Apparatus, Holliston, MA, USA) and trigger signal generator (Model 2100, A-M Systems, Sequim, WA, USA). The electrodes sat above and below the muscle to allow the length of the muscle to be imaged though the side of the bath. The proximal tendon and bone segment were secured with the attached Kevlar to the lever arm of a dual-mode servomotor (305C-LR, Aurora Scientific Inc., Aurora, ON, Canada). The jump ring attached to the distal tendon was connected to a hook either directly or via a linear extension spring. The hook was attached to a rigid post mounted onto a platform that sat on rails. The bath was filled with oxygenated amphibian Ringer’s solution of the following composition (mM): NaCl, 112; KCl, 3.3; CaCl_2_, 2.5; MgCl_2_, 1; Hepes, 20; Dextrose, 10 (pH adjusted to 7.4 with NaOH). Experiments were carried out at room temperature (21-24 °C).

### Spring properties

Maximal and submaximal force-length relationships were characterised with and without added series compliance in the form of a stainless-steel linear extension spring, giving *spring* and *no spring* conditions. In some instances, two springs were arranged in-parallel. A total of five springs were used to give seven spring configurations with the following properties: mass, 21-76 mg; stiffness, 0.10-0.70 N·mm^-1^; initial tension, 0.09-0.36 N.

### Experimental protocol

Muscle contraction was achieved with field stimulation and supramaximal stimuli of 0.1 ms duration. First, stimulation voltage was increased in 3 V increments until twitch force was maximal. The voltage giving maximum twitch force was increased further by 20%. Submaximal force was achieved with submaximal activation, specifically, twitch and doublet contractions. In most instances, twitches were unsuitable for the *spring* condition because twitch force at suboptimal lengths did not exceed the initial tension of the spring. The initial condition, *no spring* (*n* = 9) or *spring* (*n* = 8), was randomly assigned.

Each force-length relationship was characterised by performing a series of fixed-end contractions over a range of muscle lengths. Twitch and doublet (two stimuli, 100 Hz) contractions were performed first. Optimum length was determined from 8–15 contractions. Beginning with the muscle slack, muscle length was incrementally increased. A rest period of 30–60 s was given between contractions. Muscle length for tetanic contractions was assigned randomly. Tetanic contractions (100 Hz) varied in their duration because of the length dependence of force rise time and because added compliance slows the rate of force development. Stimulation duration was adjusted accordingly to give a force plateau. For the *no spring* condition, stimulation durations of 300–600 ms were sufficient at intermediate lengths, while durations up to 1 s and 1.5 s were required at the longest and shortest lengths studied, respectively. For the *spring* condition, stimulation durations typically ranged from 600 ms to 2 s. Tetanic optimum length was determined from 8–12 contractions. The rest period between tetanic contractions was at least 3–5 minutes depending on the duration of the contraction. Following each series of tetanic contractions, one or more tetanic contractions were performed at or close to optimum length to determine force loss owing to muscle fatigue and assess whether optimum length had increased during the process of constructing the force-length curve (Warren et al., 1993; Gregory et al., 2007).

Muscle sutures were video recorded at 60 Hz with a DSLR camera and macro lens that gave an image resolution of approximately 22 pixels/mm. Video recordings were calibrated with a ruler submerged in the muscle bath immediately behind the muscle. Muscle force and changes in muscle length were recorded from the servomotor and acquired at 1000 Hz using a 16-bit A/D converter (NI USB-6218, National Instruments, Austin, TX, USA). Data were displayed in Igor Pro software (WaveMetrics, Inc., Portland, OR, USA). Following the experiments, muscles were pinned in a dissection dish, extended slightly beyond slack length, and photographed for subsequent determination of pennation angle and fibre length at optimum segment length (*L*_fibre_). Muscle mass was measured after removing the distal and proximal tendons.

### Inclusion of a second no spring condition

Unexpectedly, recoil of the attached spring during relaxation for tetanic contractions led to active stretch sufficient to induce passive force enhancement and muscle damage (Fig. 1). To help characterise the damage effect, and for consistency across all 17 muscles, a second *no spring* condition was included (tetanic contractions only) when the *spring* condition was performed second, allowing for a pre-vs. post-spring comparison. In this way, maximum isometric force and optimum length were determined after spring removal for all 17 muscles. Considerations for the potential effect of stretch-induced damage on the measured difference in optimum length between *spring* and *no spring* conditions, as well as the estimate of force depression will be presented.

**Fig. 1.**
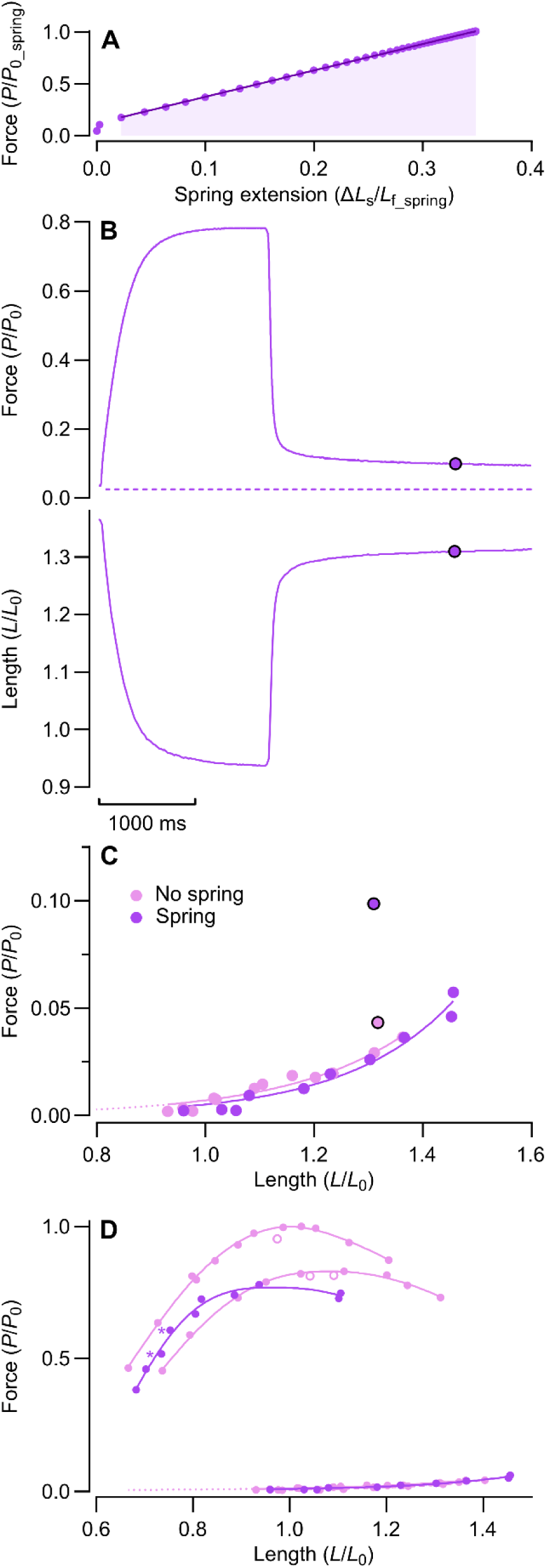
Spring recoil resulted in passive force enhancement and stretch-induced muscle damage. (A) Normalised force-extension relationship of an attached spring determined from maximum force generation. Effective spring compliance is given by normalised extension and normalised work (shaded area). Solid line represents the best fit for a linear regression fitted to data above the spring’s initial tension. (B) Force and fascicle length during a fixed-end contraction with added compliance. The attached spring was that shown in A. Circles with black outline indicate force and fascicle length 2 s after the end of stimulation. Dashed line represents predicted passive force corresponding to fascicle length 2 s after the end of stimulation. (C) Passive force-length relationship with and without added compliance. Passive force and fascicle length 2 s after the end of stimulation are plotted for contraction shown in B and for a comparable contraction without added compliance. (D) Active and passive force-length relationships for *no spring* and *spring* conditions. The *spring* condition was performed second and was followed by a second *no spring* condition. Open circles represent contractions performed at the end of a condition to assess muscle fatigue. Asterisks adjacent to active data points indicate first and second tetanic contractions performed with added compliance. *L*_0_, optimum length of initial *no sprin*g condition; *L*_0_spring_, optimum length of *spring* condition; *L*_f_spring_, fibre length at *L*_0_spring_; *P*_0_, maximum isometric force of initial no spring condition. *P*_0_spring_, maximum isometric force of *spring* condition.

### Data analysis

Data were analysed using custom-written scripts in MATLAB (The MathWorks Inc., Natick, MA, USA). High-frequency components were removed from servomotor force recordings using a zero-phase shift, second-order, 125 Hz low-pass Butterworth filter.

### Maximum isometric force and optimum length

For *spring* and *no spring* conditions, and for each contraction type, active and passive force-length relationships were constructed, and maximum isometric force and optimum length were determined. For each contraction, passive force immediately prior to stimulation and peak total force were determined, as were the corresponding passive and active segments lengths. Fascicle segment length was measured by tracking muscle sutures using a semi-automated approach from a publicly available video analysis software (Tracker, version 6.1.3, https://physlets.org.tracker). The contribution of active force to the measured total force was estimated by subtracting passive force. Because muscle shortening occurs against the stretch of series elasticity, passive force was estimated at the active fascicle length from a fitted passive force-length relationship (Scott et al., 1996; MacIntosh & MacNaughton, 2005). The passive force-length relationship at the level of the fascicle was characterised from the measurements of passive force and segment length immediately prior to the onset of stimulation. Passive force increases exponentially with length (Magid et al., 1985; Azizi & Roberts, 2010; Azizi, 2014; Meyer & Lieber, 2018), thus, passive data were fitted with an exponential equation of the form:

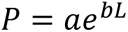

where *P* is passive force, *L* is passive fascicle length, and *a* and *b* are coefficients. The fitted passive force-length relationship was extrapolated to the minimum measured active length. For each measured active length, passive force was determined from the fitted equation and subtracted from the measured total force to give active force. Active force-length data were fitted with an equation from Otten (1987):

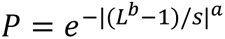

where *P* is active force, *L* is active fascicle length, and *b*, *s*, and *a* are coefficients describing the skewness, width, and roundness of the fitted curve, respectively. The fitted force-length curve was evaluated for maximum isometric force (*P*_0_) and optimum segment length (*L*_0_). Both passive and active data sets were fitted using a least-squares method.

### Evidence of active stretch during spring recoil

For contractions with added compliance, passive force following the contraction was substantially elevated compared to (1) the passive force prior to contraction, (2) the length-matched passive force derived from the fitted passive force-length relationship, and (3) the passive force following stimulation for a contraction without added compliance performed at a similar length (Fig. 1B, C). For two muscles, we confirmed that the elevated passive force after spring recoil was not a transient phenomenon. Rather, passive force remained elevated for the duration of observation, which was greater than 7 s after the end of stimulation. This passive force enhancement was evident for all muscles and interpreted as evidence of significant active stretch during spring recoil. Elevated passive force at a given length following active stretch is known as passive force enhancement and represents the passive component of stretch-induced residual force enhancement (Herzog & Leonard, 2002; Rassier et al., 2003, 2005; Herzog et al., 2003).

### Stretch-induced muscle damage

The active stretch imposed by the spring as it recoiled during force decay meant that constructing a tetanic force-length relationship with added compliance effectively exposed the muscle to a series of eccentric contractions. As few as 10 eccentric contractions can lead to an appreciable deficit in isometric force and increase in optimum length (Wood et al., 1993), although the degree of damage induced by active stretch depends on muscle length (Talbot & Morgan, 1998). Figure 1D shows a series of force-length curves for a single muscle for which contractions without added compliance were performed both before and after contractions with added compliance. Relative to the initial *no spring* curve, the final *no spring* curve was shifted rightward and exhibited force loss across the entire range of lengths studied. These features of the second *no spring* curve were consistent with stretch-induced muscle damage (Wood et al., 1993; Morgan et al., 1996; Talbot & Morgan, 1998). Contractions performed at the end of each *no spring* condition indicated only a small contribution from muscle fatigue (Fig. 1D).

### Considerations made for stretch-induced damage

Because the effective compliance given by the springs used varied several-fold, as intended, and couldn’t be easily matched across the two subsets of muscles, added compliance was treated as a continuous variable rather than a dichotomous one (i.e., *spring* or *no spring*). On this basis, we opted to compare the *spring* curve to the following *no spring* curve for all 17 muscles, and perform additional analyses to determine to what extent, if at all, stretch-induced damage may have confounded our findings. This comparison was further justified because it did not appear to be influenced by an initial *no spring* condition. When effective spring compliance was similar for two muscles, nearly identical force-length curves for the *spring* condition were obtained when normalising to the *no spring* condition performed after spring removal (Fig. 2). The effect of a matched amount of added compliance on force depression and *L*_0_, when evaluated in this manner, did not appear to depend on whether a series of *no spring* contractions was performed first. Thus, it was possible to examine the relationship between effective spring compliance and force depression, or optimum length, for all 17 muscles using a single regression, treating added compliance as a continuous variable.

**Fig. 2.**
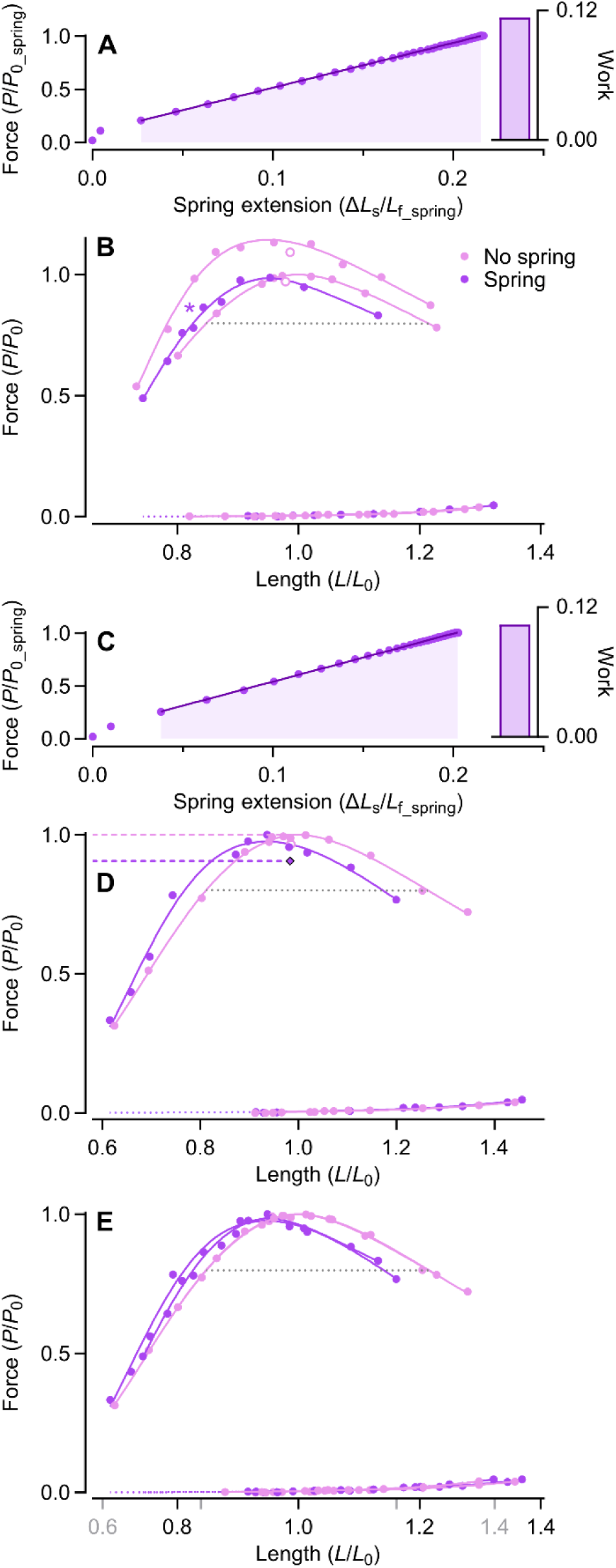
Justification for assessing the effect of added compliance by comparing *spring* data to following *no spring* data despite an initial no *spring* condition. (A, C) Normalised force-extension relationship of two springs giving similar effective compliances. *Inset*, normalised work, which is given by the shaded area. (B, D) Force-length relationship with and without added compliance for the two muscles corresponding to A and C, respectively. In *B*, the *spring* condition was performed second. Data are normalised to *P*_0_ and *L*_0_ of the *no spring* condition that followed the *spring* condition. Open circles represent a contraction performed at the end of the condition to assess muscle fatigue. Asterisk adjacent to active data point indicates first tetanic contraction performed with added compliance. In *D*, the *spring* condition was performed first. Data are normalised to *P*_0_ and *L*_0_ of the following *no spring* condition. Dotted grey lines represent width at 0.8 *P*_0_. Diamond with outline represents active force from a final contraction performed with added compliance. The difference in force between this final contraction and *P*_0_ for the following *no spring* condition (dashed lines) gave a second estimate of force depression that accounted for the loss of force due to stretch-induced muscle damage. (E) Force-length relationship from the *spring* condition and subsequent *no spring* condition for both muscles are superimposed. The x-axis scale for the first muscle (grey) has been adjusted such that the curve widths at 0.8 *P*_0_ for the *no spring* curves are matched.

The shift in *L*_0_ and reduction in *P*_0_ between two *no spring* conditions (*n* = 9) were used as metrics of stretch-induced muscle damage (Wood et al., 1993; Talbot & Morgan, 1998; Whitehead et al., 2003). The association between these metrics and effective spring compliance was determined to confirm that our findings weren’t confounded by stretch-induced damage. Because stretch-induced damage would lower any measurement of force after spring removal, a second estimate of force depression was determined for a subset of muscles (*n* = 12) by taking maximum force with added compliance from a final contraction performed at or close to *L*_0_ before removing the spring (Fig. 2D, Fig. 3E). In this way, stretch-induced damage equally affected the measured maximum force with and without added compliance such that its contribution to the estimate of force depression was minimised.

**Fig. 3.**
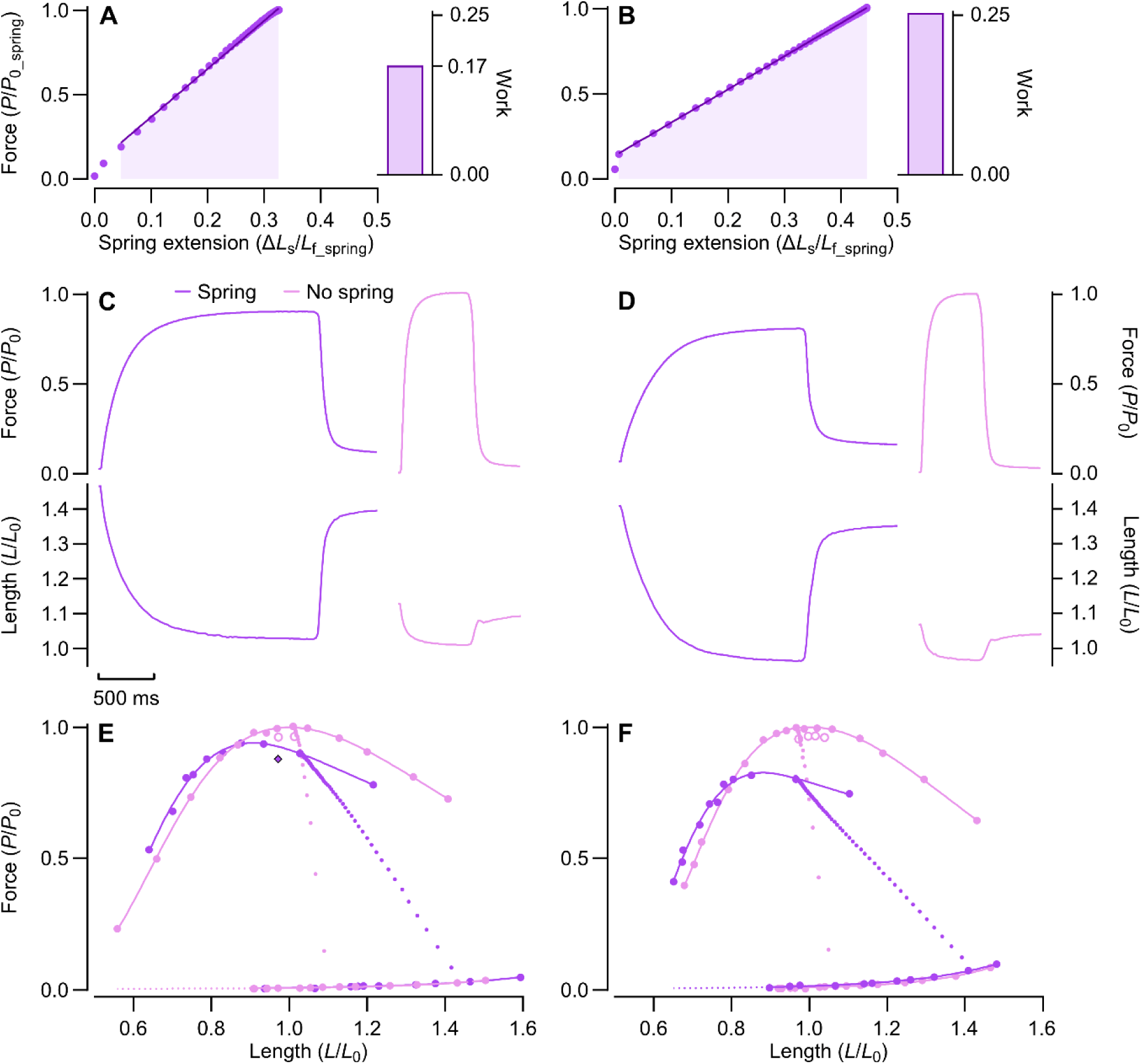
Force and fascicle length during fixed-end contractions with and without added compliance and associated force-length relationship. (A, B) Normalised force-extension relationship of two springs with different effective compliances. *Inset*, normalised work, which is given by the shaded area. (C, D) Force and fascicle length during a fixed-end contraction with and without added compliance. The isometric fascicle length for each contraction was ∼1.0 *L*_0_. (E, F) Active and passive force-length relationships with and without added compliance. For both muscles, the *spring* condition was performed first. Dotted lines connecting passive and active curves represent force-length trajectories for contractions shown in C and D. Open circles represent contractions performed at the end of the condition to assess muscle fatigue. Diamond with outline in E represents active force from the final contraction of the *spring* condition.

### Force depression and shift in optimum length

In the following text, *P*_0_ and *L*_0_ notation are reserved for the *no spring* condition following spring removal, while *P*_0_spring_ and *L*_0_spring_ notation are used for the *spring* condition. Both sets of notations are used exclusively for tetanic contractions. To evaluate the relationship between effective spring compliance and force depression, or optimum length, *P*_0_spring_ was expressed relative to *P*_0_, and *L*_0_spring_ was expressed relative to *L*_0_. Force depression was defined as a reduction in *P*_0_spring_ with respect to *P*_0_. This comparison was preferred with respect to a traditional comparison of force at a matched steady-state length because it represents the deficit in maximum tetanic force, and we predicted that force depression would alter the shape of the force-length relationship.

### Effective spring compliance

The effective compliance of an attached spring or springs was determined for the tetanic contraction giving maximum force. Spring extension was measured by tracking the spring-muscle interface using the methods described for tracking muscle sutures. Because force depression increases as a function of mechanical work performed during shortening (Granzier & Pollack, 1989; Herzog et al., 2000; Van Noten & Van Leemputte, 2011), we expressed effective spring compliance as the maximum work performed by the muscle on the attached spring. Muscle work was determined from the area under the extension-force curve when force was greater than the initial tension of the spring. Muscle work was normalised to the force-generating capacity (*P*_0_spring_) and optimum fibre length of the muscle when shortening against the spring (*L*_f_spring_), which gave a dimensionless measure of work (Kosterina et al., 2009). *L*_f_spring_ was given by scaling relative segment length (i.e., *L*_0_spring_/*L*_0_) according to the group average for *L*_f_, which gave stronger correlations than individual muscle *L*_f_. The capacity to perform work against a spring depends only on the maximum force that can applied to the spring. Since the amplitude of shortening is frequently reported in force depression literature, effective spring compliance was also expressed as maximum spring extension normalised to the group average for *L*_f_ [i.e., Δ*L*_s_/*L*_f_ (Biewener & Roberts, 2000)]. These two representations of effective spring compliance, normalised work and normalised spring extension, were very strongly related (*P* < 0.001, *r*^2^ = 0.99), as expected. Since the spring was mechanically in-series with the muscle, we retained both representations as normalised spring extension can be directly related to the ‘fixed-end compliance’ (FEC), defined as fibre shortening against the stretch of series elasticity normalised to optimum fibre length (Roberts, 2002).

### Activation dependence of optimum length

The effect of added compliance and shortening-induced force depression on the activation dependence of optimum length was assessed in two ways. First, we evaluated the relationship between effective spring compliance and the shift in *L*_0_spring_ relative to *L*_0_. Second, we evaluated the relationship between effective spring compliance and the difference between *L*_0_spring_ and the optimum length for a doublet contraction with added compliance. The activation-related shift in optimum length was expressed as a percentage of *L*_0_spring_. Twitch and doublet contractions without added compliance were compared against *L*_0_.

### Muscle contractile and morphological properties

The tetanic contraction that generated the greatest active force (*F*_max_) within each respective condition was evaluated for 50 and 90% rise time. *F*_max_ was expressed relative to muscle physiological cross-sectional area (PCSA) to determine maximum isometric stress. PCSA was calculated according to the equation of Sacks and Roy (1982):

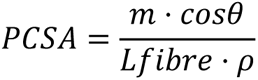

where *m* is muscle mass, *θ* is fibre pennation angle, and *ρ* is muscle density. Muscle density was assumed to be 1.06 g·cm^-3^ (Leonard et al., 2021). *L*_fibre_ was determined from 3-5 measurements of the curved trajectory between fascicle endpoints using ImageJ software (National Institutes of Health, Bethesda, MD, USA). Measurements were made from superficial fascicles passing through the mid-belly of the muscle. The average of the measured lengths was scaled according to the relative length of the fascicle segment (*L*/*L*_0_) at the pinned muscle length to give *L*_fibre_, fibre length at *L*_0_. Pennation angle was measured for each measured fascicle length and averaged.

### Statistics

A least-squares linear regression was used to characterise the relationship between effective spring compliance (i.e., normalised work, normalised spring extension) and the dependent variable of interest, including force depression (i.e., *P*_0_spring_), shift in tetanic optimum length (i.e., *L*_0_spring_), and activation-related shift in optimum length. A one sample *t* test was used to determine whether the shift in optimum length for submaximal contractions differed from a hypothetical value of 0. Fitted regression lines are plotted with the 95% confidence interval (CI). Group data are displayed as the mean ± 95 CI or as box and whisker plots, where the box extends to the first and third quartiles and the whiskers extend to the minimum and maximum values. Unless stated otherwise, data are reported as the mean ± the standard deviation (SD). Statistical significance was set at *P* < 0.05. Statistical analyses and data visualisation were performed using Prism software (GraphPad Software Inc., La Jolla, CA, USA).

## RESULTS

Morphological and contractile properties for the 17 bullfrog semitendinosus muscles studied to determine the effect of added compliance on tetanic optimum length and the activation dependence of optimum length are reported in Table 1.

**TABLE 1.**
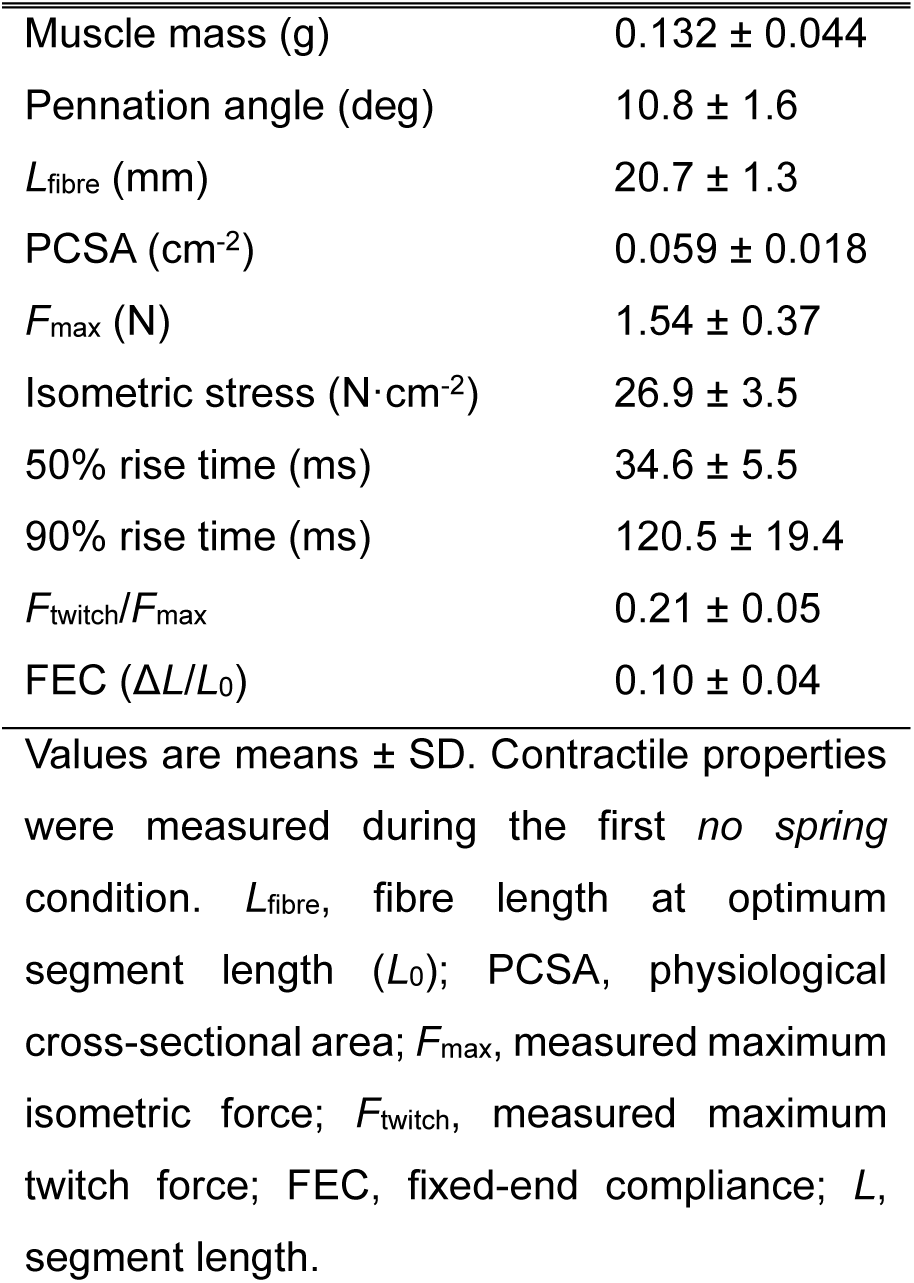
Morphological and contractile properties of the bullfrog semitendinosus muscle (dorsal head)

|  |  |
| --- | --- |
| Muscle mass (g) | $0.132 \pm 0.044$ |
| Pennation angle (deg) | $10.8 \pm 1.6$ |
| $L_{\text{fibre}}$ (mm) | $20.7 \pm 1.3$ |
| PCSA ($\text{cm}^2$) | $0.059 \pm 0.018$ |
| $F_{\text{max}}$ (N) | $1.54 \pm 0.37$ |
| Isometric stress ($\text{N} \cdot \text{cm}^{-2}$) | $26.9 \pm 3.5$ |
| 50% rise time (ms) | $34.6 \pm 5.5$ |
| 90% rise time (ms) | $120.5 \pm 19.4$ |
| $F_{\text{twitch}}/F_{\text{max}}$ | $0.21 \pm 0.05$ |
| FEC ($\Delta L/L_0$) | $0.10 \pm 0.04$ |

Values are means ± SD. Contractile properties were measured during the first *no spring* condition. $L_{\text{fibre}}$ , fibre length at optimum segment length ( $L_0$ ); PCSA, physiological cross-sectional area; $F_{\text{max}}$ , measured maximum isometric force; $F_{\text{twitch}}$ , measured maximum twitch force; FEC, fixed-end compliance; $L$ , segment length.

### Effects of added compliance on active shortening, force generation, and the force-length relationship

Figure 3 shows recordings of force and fascicle length during representative fixed-end contractions with and without added compliance for two muscles. Effective spring compliance, illustrated as normalised work and normalised spring extension (i.e., FEC), was noticeably different for the two muscles (Fig. 3A, B). For both muscles, steady-state force was lower for the *spring* compared to *no spring* contraction despite similar steady-state fascicle lengths. Force-length curves constructed from tetanic contractions with and without added compliance show that maximum isometric force was lower with added compliance, whilst optimum length was shorter (Fig. 3E, F). Compared to the *no spring* condition, the force-length curve with added compliance was leftward-shifted and depressed such that there was a force deficit across the plateau region and on the descending limb. Because the *spring* condition preceded the *no spring* condition, the depression of force was not owing to muscle fatigue or stretch-induced damage. Force loss owing to the latter, however, was likely responsible for the apparent absence of force depression on the ascending limb. Force depression on the ascending limb was confirmed after evaluating instances where the first or second tetanic contraction with added compliance fell on the ascending limb and could be compared against an initial *no spring* condition (Fig. 1D and Fig. 2B).

### Effective spring compliance as a determinant of force depression and optimum length

Force depression, which we defined as a deficit in *P*_0_spring_ relative to *P*_0_, increased as a function of the effective compliance added. Specifically, a very strong, negative correlation existed for the relationship between normalised work and *P*_0_spring_ normalised to *P*_0_ (*P* < 0.001, *r*^2^ = 0.89; Fig. 4A). An equally strong relationship existed when effective compliance was represented as normalised spring extension (*P* < 0.001, *r*^2^ = 0.91; Fig. 4A, Inset). Added compliance also reduced tetanic optimum length in a compliance-dependent manner. The relationship between muscle work performed on the spring and *L*_0_spring_ normalised to *L*_0_ was well fitted by a negative linear regression (*P* < 0.001, *r^2^* = 0.63; Fig. 4B). Representing effective spring compliance as normalised spring extension gave an equally strong correlation (*P* < 0.001, *r^2^* = 0.60; Fig. 4B, Inset).

**Fig. 4.**
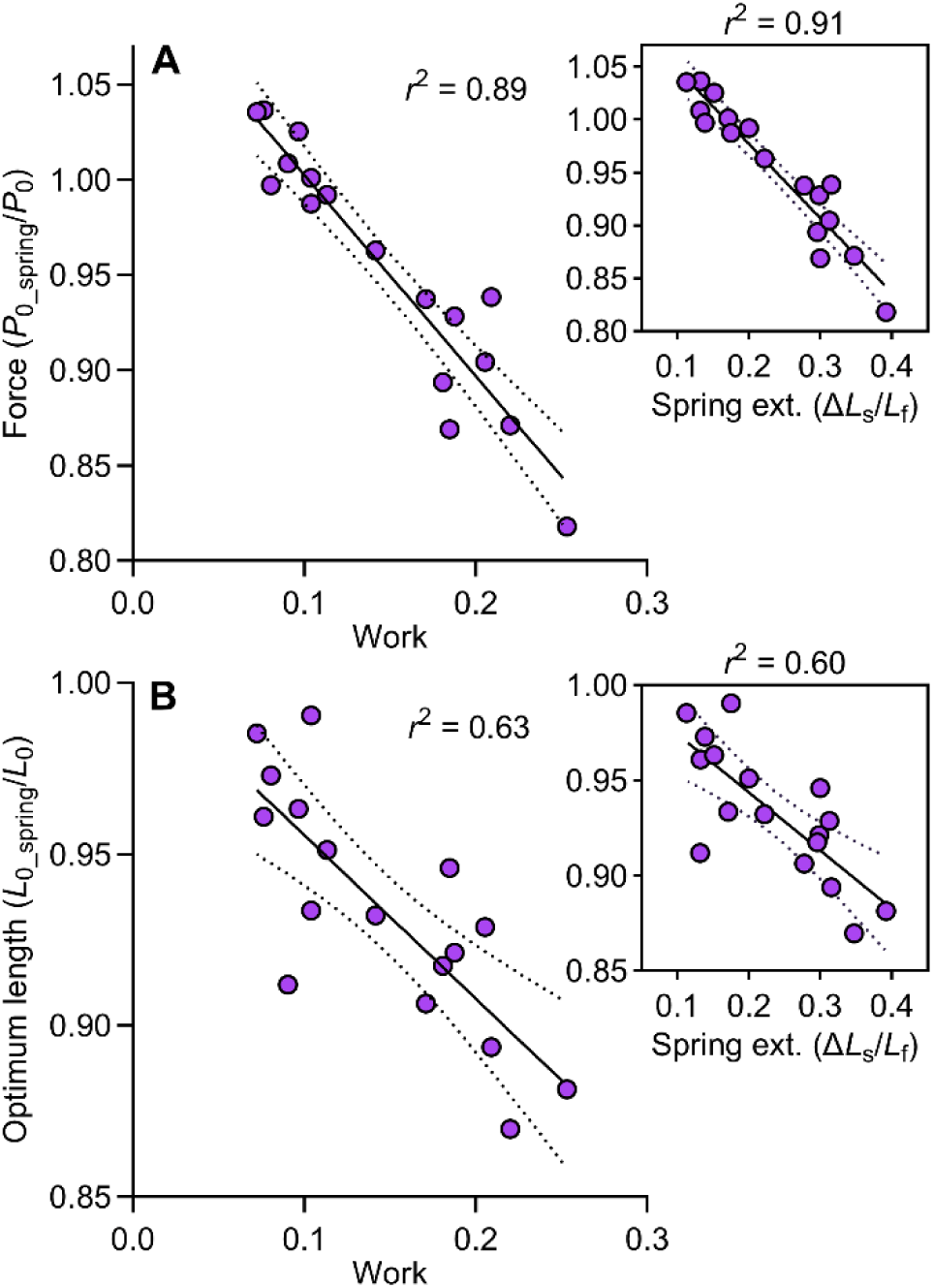
Mechanical work and spring extension as determinants of force depression and shift in optimum length. (A) Relationship between effective spring compliance, defined as normalised work performed on the spring, and maximum tetanic force (*P*_0_spring_) normalised to *P*_0_, which we used as an index of force depression (*n* = 17). *Inset*, effective compliance represented as normalised spring extension. (B) Relationship between normalised work and tetanic optimum length (*L*_0_spring_) normalised to *L*_0_ (*n* = 17). *Inset*, effective spring compliance represented as normalised spring extension. *L*_s_, spring length; *L*_f_, optimal fibre length.

### Activation dependence of optimum length

The activation dependence of optimum length was assessed using twitch and doublet contractions (Fig. 5A). For the brief doublet contraction, added compliance slowed force development such that peak force was similar to twitch force without added compliance (Fig. 5B), giving a force-similar comparison in addition to an activation-matched comparison. Maximal and submaximal force-length relationships with and without added compliance for a representative muscle are shown in Figure 5C. The activation dependence of optimum length with added compliance was positively correlated with muscle work performed on the spring (*P* = 0.0117, *r*^2^ = 0.45; Fig. 5D). In the absence of added compliance, the activation dependence of optimum length was modest. Doublet optimum length (0.99 *L*_0_, 95% CI: 0.98, 1.01) did not differ with respect to tetanic optimum length (*P* = 0.274, η_p_^2^ = 0.09; *n* = 14), whilst twitch optimum length (1.03 *L*_0_, 95% CI: 1.02, 1.05) was just 3.2% longer than tetanic optimum length (*P* < 0.001, η_p_^2^ = 0.56; *n* = 17).

**Fig. 5.**
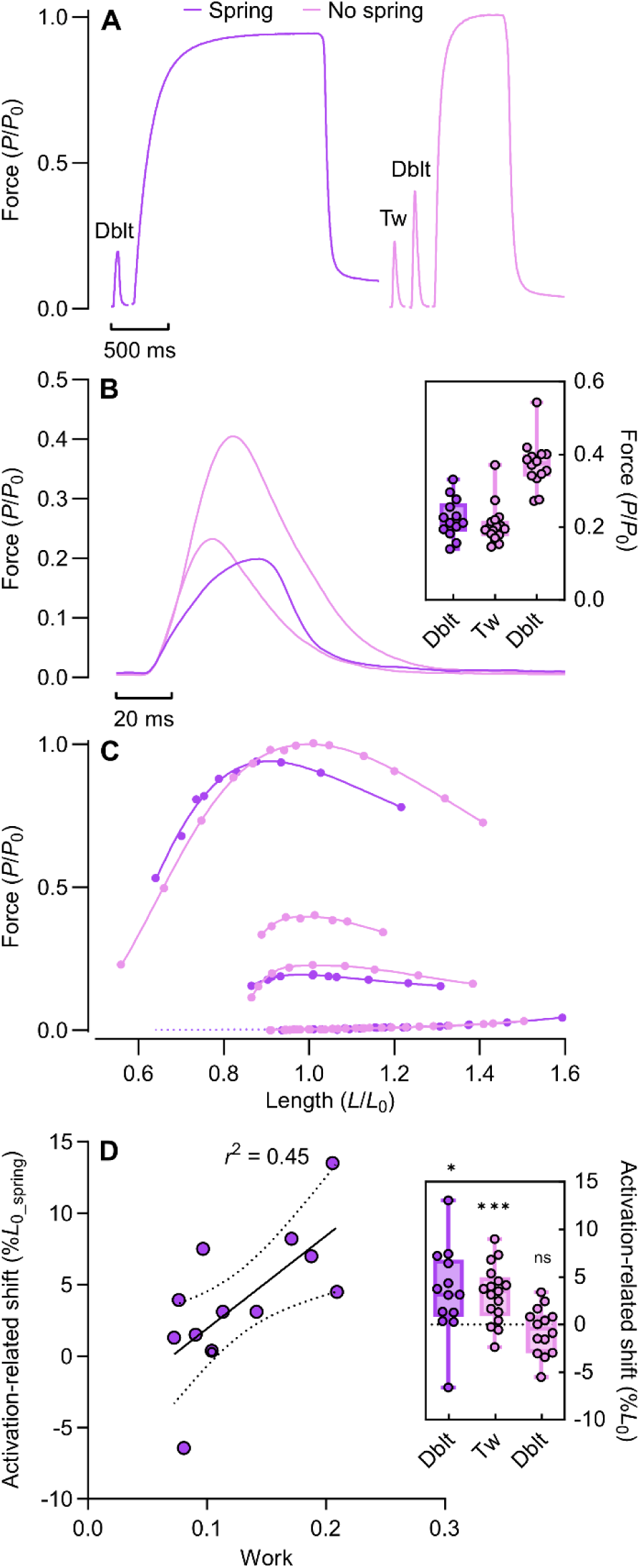
Activation dependence of optimum length. (A) The effect of submaximal activation on optimum length was assessed using twitch (*no spring*) and doublet (*no spring*, *spring*) contractions. (B) Time course of force for submaximal contractions with and without added compliance. *Inset*, maximum force for twitch and doublet contractions. (C) Maximal and submaximal force length relationships with and without added compliance for a representative muscle. (D) Relationship between effective spring compliance and activation-related shift in optimum length (*n* = 13). *Inset*, Activation-related shift in optimum length with and without added compliance (Twitch: *n* = 17; Doublet: *n* = 14). One sample *t* test (hypothetical value = 0): *, *P* = 0.020; ***, *P* < 0.001; ns, *P* = 0.342.

### Stretch-induced muscle damage

Stretch-induced muscle damage manifested as additional force loss compared to that evident after a series of fixed-end contractions without added compliance (Fig. 1D, Fig.2B and Fig. 3E, F), and an increase in optimum length between two *no spring* conditions (Fig. 1D, 2B). This force loss force likely lowered the measured force depression. For the lowest effective spring compliances, *P*_0_spring_ exceeded *P*_0_, and the y-intercept of the linear regression fitted to the data was 1.11 *P*_0_ (95% CI: 1.08, 1.14; Fig. 4A). Presumably, this result reflects an underestimation of force depression, which can be attributed to an underestimation of *P*_0_ after the spring was removed because of stretch-induced damage and fatigue. We attempted to account for the component of stretch-induced damage by taking the force generated by a final contraction close to *L*_0_spring_ before removing the spring. Using this analysis for a subset of muscles (*n* = 12), force depression remained strongly correlated with effective compliance (*P* < 0.001, *r*^2^ = 0.89) but was greater for a given effective compliance such that force depression was now evident for the stiffest springs (Fig. 6A). The y-intercept of this regression, –4.1% (95% CI: –0.4, –7.8), was taken as an estimate of muscle fatigue and agreed with our measurements of force loss evident after constructing a single *no spring* force-length curve (before *spring*: 0.05 *P*_0_; after *spring*: 0.02–0.03 *P*_0_). The similarity in slope for the two quantifications of force depression indicated that damage was largely independent of the amount of added compliance.

**Fig. 6.**
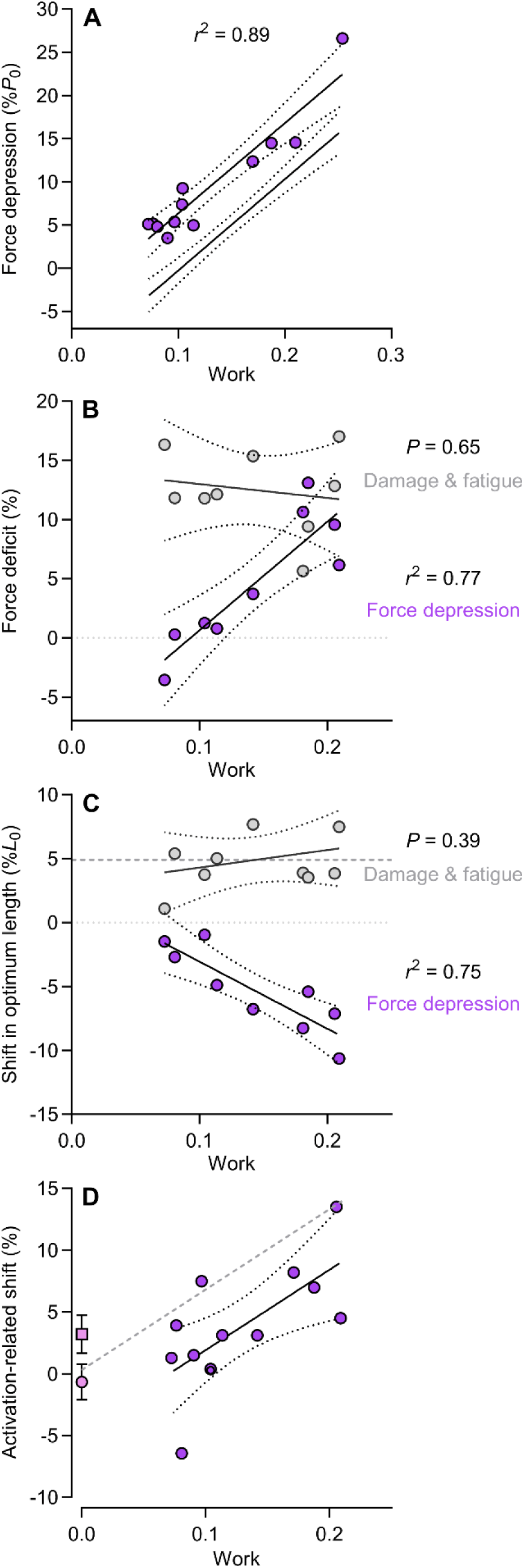
Effects of stretch-induced damage on force depression and shift in optimum length. (A) Relationship between effective spring compliance and force depression after accounting for stretch-induced damage (*n* = 12). Also shown is the fitted regression line from the original analysis of all 17 muscles originally shown in Fig. 4A. (B) Relationship between effective spring compliance and the force deficit evident between *no spring* curves (i.e., pre-spring and post-spring) due to damage and fatigue. Also shown is the relationship between effective spring compliance and the deficit in maximum isometric force with added compliance (i.e., force depression) for the same subset of muscles (*n* = 9). (C) Relationship between effective spring compliance and shift in optimum length evident between *no spring* curves (i.e., pre-spring and post-spring) due to damage (and possibly fatigue). Also shown is the relationship between effective spring compliance and the shift in optimum length with added compliance attributed to force depression for the same subset of muscles (*n* = 9). Dashed grey line at 4.9% *L*_0_initial_ represents the average shift. Neither index of damage was associated with effective spring compliance. (D) Comparison of the activation dependence of optimum length with and without added compliance. Data at 0 work represent the shift measured for twitch (square) and doublet (circle) contractions without added compliance. The relationship between effective spring compliance and activation dependence of optimum length is shown in its original form and with an offset of 4.9% *L*_0_ applied to the fitted regression (grey dashed line). This offset is equal to the average shift in tetanic optimum length attributed to stretch-induced damage (see B).

The confounding effect of stretch-induced damage was tested by determining whether the loss of maximum force and shift in optimum length (4.9%, *P* < 0.001, η ^2^ = 0.84) evident between *no spring* curves (i.e., before vs after *spring*) were related to effective spring compliance (Fig. 6B, C). We found that there was no correlation between muscle work and either the extent to which maximum force had decreased (*P* = 0.651, *n* = 9; Fig. 6B) or the extent to which optimum length had increased (*P* = 0.397; *n* = 9; Fig. 6C) after the added compliance was removed. For this same subset of muscles, force depression and optimum length with added compliance were positively correlated with effective spring compliance (*P* = 0.002, *r*^2^ = 0.75-0.77; *n* = 9; Fig. 6B, C), as was the case when all 17 muscles were included in the regression.

It appeared that the effect of stretch-induced damage on optimum length was realised during the measurement of optimum length with added compliance. The y-intercept of the regression for the relationship between effective spring compliance and shift in optimum length was 1.003 *L*_0_ (95% CI: 0.97-1.04; Fig. 4B), while the y-intercept for the relationship between added compliance and the activation dependence of optimum length was –4.6% (95% CI: –11.0, 1.9; Fig. 5D). This value represents a deviation from what we anticipated since doublet optimum length did not differ with respect to tetanic optimum length in the absence of added compliance (Fig. 5D, Inset) and added compliance lowered peak doublet force considerably (Fig. 5B, Inset). The negative y-intercept presumably reflects a damage-induced increase in tetanic optimum length with (and without) added compliance, as doublet optimum length was always measured before tetanic optimum length. Applying an offset to the relationship between added compliance and the activation-related shift of optimum length equal to the shift in tetanic optimum length attributed to stretch-indued damage (i.e., 4.9% *L*_0_, see Fig. 6C) gave a new y-intercept of 0.3% (i.e., 1.003 *L*_0_), which fell between the shifts measured without added compliance for the force-similar twitch and activation-matched doublet (Fig. 6D).

## DISCUSSION

The activation dependence of optimum length isn’t satisfactorily explained by the length dependence of Ca^2+^ sensitivity (Holt & Azizi, 2014; Holt & Williams, 2018; Hessel et al., 2019). Because active shortening occurs against the stretch of series elasticity during force development, and muscle preparations of differing compliance differ in their activation dependence of optimum length (Holt & Williams, 2018), shortening-induced force depression has been identified as a potential determinant of optimum length. Critically, force depression is work– and length-dependent, and the interaction of these dependencies provides another means for the force-length relationship to be uncoupled from myofilament overlap. The *force depression hypothesis* proposes that for a muscle with significant series compliance, the optimum length for a tetanic contraction (i.e., high force, high work) is shorter than the length corresponding to optimal filament overlap (Holt & Williams, 2018). For a low-force contraction, optimum length should be relatively unaffected by series compliance because force depression is proportional to muscle work, and work performed on a spring is proportional to active shortening squared. A work-dependent modulation of optimum length would mean that the activation dependence of optimum length would increase as series compliance and work during force development increase.

We tested these predictions by adding different amounts of in-series compliance to isolated muscles and constructing force-length relationships for maximal and submaximal contractions. In agreement with the *force depression hypothesis*, optimum length for a tetanic contraction decreased in proportion to both the effective compliance of an attached spring and the associated force depression, while the activation dependence of optimum length increased in proportion to effective spring compliance. Together, these findings implicate force depression as a determinant of the activation or force dependence of optimum length.

### Series compliance, force depression, and the activation dependence of optimum length

According to the relationship between effective spring compliance and tetanic optimum length, for every 0.10 *L*_f_ increase in fascicle shortening during maximum force generation, optimum length decreased by 0.031 *L*_0_. We believe this finding to be the first direct evidence that the activation or force dependence of optimum length could arise in part from tetanic optimum length being shorter than the length that presumably corresponds to optimum filament overlap. We propose that this reduction in optimum length with added compliance can be attributed to shortening-induced force depression. Force depression increases in proportion to work (Herzog et al., 2000; Corr & Herzog, 2005; Van Noten & Van Leemputte, 2011) but also increases exponentially as a function of muscle length (Van Noten & Van Leemputte, 2011). The interaction of these factors with the length dependence of myofilament overlap (i.e., the potential to generate force and, therefore, do work), would appear to provide the right conditions for added compliance to decouple maximum force generation from optimum filament overlap. In agreement with conventional experiments of shortening-induced force depression, we found that force depression was very strongly correlated to both muscle work and fascicle shortening amplitude.

The contribution of series compliance and force depression to the activation dependence of optimum length was previously ruled out after a similar experiment on the rat medial gastrocnemius muscle using silicone tubing to increase active shortening (MacDougall et al., 2020). However, our findings indicate that the absence of a compliance-mediated reduction in optimum length in this prior work could simply reflect insufficient added compliance and relatively low activation such that the mechanical work threshold for appreciable force depression was never reached. During the triplet contraction, which generates a submaximal force, the added compliance given by the silicone tubing increased active shortening by less than 0.02 *L*_0_. In contrast, we increased active shortening by 0.24 *L*_0_ on average (range: 0.11-0.39 *L*_0_) using a linear extension spring of prescribed compliance and maximal activation. According to our regression equations for the relationship between effective spring compliance, force depression, and shift in optimal length, the compliance added by the silicone tubing would have induced a 0.01 *P*_0_ deficit in force and reduced optimum length by 0.006 *L*_0_. It’s unlikely that any method for determining the force-length relationship of skeletal muscle would possess the level of precision required to detect such a small shift in optimum length.

The sensitivity of force depression to work during shortening is a necessary element if force depression is to mediate an increase in the activation dependence of optimum length. Although we could not quantify force depression for the doublet since force generation is only transient and attenuated due to force-velocity effects, we did demonstrate that the activation dependence of optimum length was positively correlated with spring compliance. We can infer from this finding that added compliance had a comparatively smaller effect on optimum length for our low-force contraction. We predict that the actual shift might be very modest since the work performed against a spring increases in proportion to displacement squared, which means that work and force depression diminish more rapidly when activation and force are lowered. We considered the most compliant spring added, for which maximum extension represented 0.39 *L*_0_. In terms of fixed-end compliance, this value likely approaches the upper limit of the physiological range. If force were to be lowered to 0.25 *P*_0_spring_, which is similar to doublet force with added compliance, work done would be 1/16 of tetanic work, and force depression would fall from 27% to just 2%. The resulting shift in optimum length would be anticipated at just 0.008 *L*_0_. In contrast, tetanic optimum length would be reduced by 0.12 *L*_0_ for this level of added compliance and, therefore, force depression alone would produce an activation dependence of optimum length of approximately 11%. This analysis also reveals that the deficit in force evident for doublet contractions with added compliance (0.22 *P*_0_ vs 0.37 *P*_0_) was due almost entirely to an increased speed of shortening speed and slower force development.

Indirect support for the involvement of series compliance and force depression in determining optimum length previously came from the observed difference in the activation dependence of optimum length between a bullfrog plantaris fibre bundle and intact bullfrog plantaris muscle. Holt and Williams (2018) found that the difference in optimum length between twitch and tetanic contractions was 35% for the intact muscle but just 8% for the fibre bundle. The 8% increase observed for the fibre bundle, a preparation with little series compliance, especially relative to the intact muscle, was attributed to the length dependence of Ca^2+^ sensitivity. The difference in the activation dependence between the two preparations, roughly 0.27 *L*_0_, was attributed to differences in series compliance, namely the potential for force depression to arise from active shortening against the stretch of tendon and aponeurosis.

Our findings indicate that force depression likely contributes to, but does not fully explain, differences in the activation dependence of optimum length evident between muscle preparations with vastly different compliances. The fixed-end compliance of the bullfrog plantaris muscle is approximately 30% (Sawicki et al., 2015). For this muscle, we would predict that force depression would cause tetanic optimum length to be reduced by approximately 0.09 *L*_0_ compared to the length corresponding to optimum filament overlap. This shift falls short of the 0.27 *L*_0_ shift predicted from the comparison of fibre bundle and intact muscle preparations. Some of this discrepancy could be due to muscle-specific work and length dependencies of force depression. Alternatively, the effect of series compliance might have been previously overestimated. The explanatory power of the work comparing the activation dependence of optimum length between bullfrog plantaris preparations is somewhat limited because optimum fibre lengths could not be directly compared. It remains possible that the apparent effect of series compliance demonstrated by the referenced study was the result of a more complex structural disparity, possibly related to intramuscular connective tissues, internal work, and force transmission (Holt & Azizi, 2014).

It is widely accepted that the activation dependence of optimum length exhibited by whole muscle is owing to length-dependent Ca^2+^ sensitivity (Rassier et al., 1999; MacIntosh, 2017), a property of the contractile apparatus. Corresponding measurements of the force-Ca^2+^ and force-length relationships in single fibres show that the elevated Ca^2+^ sensitivity at long lengths is sufficient to shift optimum length beyond optimal filament overlap when activation is submaximal (Balnave and Allen, 1996; de Beer et al., 1988; Moss et al., 1983; Stienen et al., 1985). The magnitude of the activation-related shift at this structural level is generally 15-30% (Balnave and Allen, 1996; de Beer et al., 1988; Moss et al., 1983; Stienen et al., 1985; Zuurbier et al., 1998, 1995). We found that the activation dependence of optimum length for the bullfrog semitendinosus muscle was modest. Optimum length for a twitch (without added compliance) was just 3% longer than *L*_0_, and there was no shift in doublet optimum length despite a relatively low force (0.37 *P*_0_). Although a relatively small shift in optimum length could indicate a compliant titin isoform (Fukuda et al., 2003; Patel et al., 2012; Mateja et al., 2013), growing evidence suggests that the activation dependence of optimum isn’t solely determined by the length dependence of Ca^2+^ sensitivity (Holt & Azizi, 2014; Holt & Williams, 2018; Hessel et al., 2019). In fact, Holt and Azizi (2014) provided evidence of a strong Ca^2+^-independent mechanism. They found that optimum length increased dramatically as muscle recruitment was decreased, showing a dependence on contraction force rather than thin filament activation. Our findings illustrate that shortening-induced force depression could be partly responsible for this apparent force dependence (Holt & Williams, 2018).

We considered the possibility that the pronounced activation dependence exhibited by the bullfrog plantaris muscle could be explained further by the interactive effect of compliance and length on force rise time. Series compliance slows and prolongs force development (Fig. 7A, B), while the relationship between length and rise time is ‘U’ shaped with a minimum at lengths shorter than optimum length (Fig. 7C). Between 0.8 and 1.2 *L*_0_, 90% rise time with added compliance increased three-fold (Fig. 7D). If the duration of stimulation selected for a particular study was insufficient for true steady-state force, the underestimation of force would correspond to the length-dependence of force rise time. Thus, brief stimulation durations could cause the force-length relationship to be shifted leftward with respect to filament overlap. We simulated the use of a briefer stimulation duration by taking peak force and the corresponding fascicle length over the initial 300 ms after the onset of force development (Fig. 7E). As predicted, this reanalysis produced a systematic reduction in tetanic optimum length for the added compliance condition (range: 0.01–0.06 *L*_0_spring_, Fig. 7E, inset). However, since the average reduction was just 0.03 *L*_0_spring_, it seems unlikely that stimulation duration is a critical factor determining the variation in the activation dependence of optimum length. The predominant factor contributing to the the large force dependence of optimum length for the bullfrog plantaris muscle remains elusive. Although the findings from motor unit recruitment studies are generally inconsistent with the internal work-force transmission hypothesis, we can’t rule out the possibility that the unique behaviour of the plantaris muscle is owing to muscle-specific properties.

**Fig. 7.**
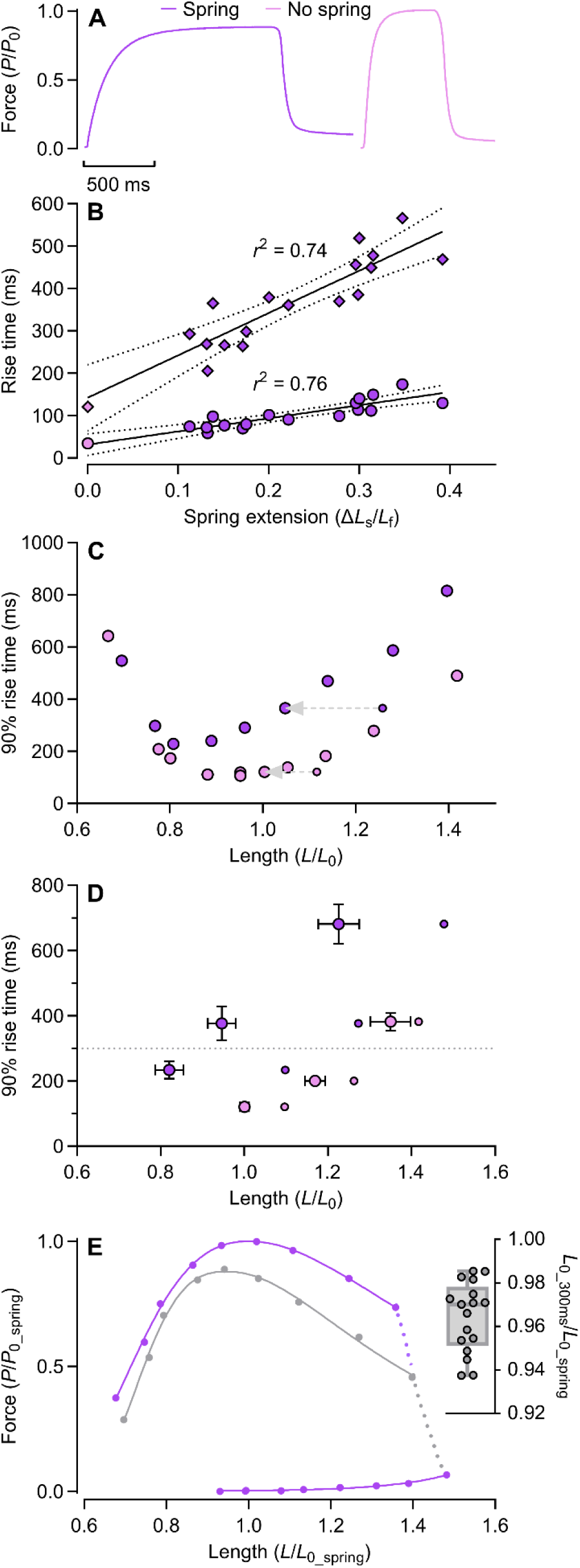
Force rise time as a function of added compliance and fascicle length. (A) Time course of force rise for the tetanic contraction giving maximum force with (left) and without (right) added compliance. (B) Relationship between maximum work performed on spring and 50 and 90% rise times for the tetanic contraction giving maximum force. Data points at 0 work are means and 95% CIs for tetanic rise times measured without added compliance. (C) 90% rise time as a function of active fascicle length with and without added compliance for the muscle corresponding to panel A. Small circles and grey arrows illustrate the initial fascicle length and shortening magnitude for the contractions giving maximum force. (D) Mean 90% rise time and 95 CI as a function of active fascicle length for three matched initial fascicle lengths corresponding to (1), optimal initial length without added compliance, (2) optimal initial length with added compliance, and (3) longest initial length. € Tetanic force-length relationship measured with added compliance for the muscle corresponding to panels A and C. The lower fitted curve is constructed from the peak force and corresponding fascicle length measured within the first 300 ms after the onset of stimulation for each contraction. Abbreviating the time allowed for peak force to develop reduced tetanic optimum length (*L*_0_300ms_) by 5.4% for this muscle. Inset, *L*_0_300ms_ for each muscle.

### Involvement of stretch-induced muscle damage

An unexpected consequence of using linear extension springs as added compliance was that spring recoil during force decay caused sufficient active stretch to induce passive force enhancement and muscle damage. We found that tetanic optimum length was longer once the spring was removed compared to before spring attachment. This finding could be used to argue that the difference in optimum length with and without compliance was simply due to stretch-induced damage rather than force depression. However, the shift in optimum length evident between *no spring* conditions was independent of effective compliance (Fig. 6C). Moreover, two results lead us to believe that the increase in optimum length resulting from stretch-induced damage was realised during the measurement of optimum length with added compliance (Fig. 4B, Fig.5D, and Fig. 6C, D), in which case, we can attribute the apparent reduction in tetanic optimum length mediated by added compliance to force depression.

If we accept that the increase in optimum length resulting from damage was largely accounted for during the measurement of optimum length with added compliance, the difference in force at any given length with and without added compliance must largely reflect the net effect of force depression and subsequent force loss due to damage and fatigue. We believe the pattern of force depression observed is generally consistent with the length dependence of force depression (Edman et al., 1993; Meijer et al., 1998; Morgan et al., 2000; Van Noten & Van Leemputte, 2011). Given that we were able to confirm the presence of some force depression on the ascending limb (Fig. 1D and Fig. 2B), we propose that relatively little force depression developed at lengths on the ascending limb such that the subsequent loss of force due to damage resulted in a net deficit in force when added compliance was removed. At intermediate and long lengths, force depression likely exceeded force loss since there was a net gain in force when added compliance was removed. This apparent length dependence of force depression is required for added compliance, via force depression, to reduce tetanic optimum length. A work-dependence alone would result in a downward shift of the force-length relationship only. Since stretch-induced damage cannot account for our findings, and we have shown that the force depression for fixed-end contractions is both work– and length-dependent, we must conclude that force depression was primarily responsible for the observed relationships between effective spring compliance, tetanic optimum length, and the activation dependence of optimum length.

### Considerations for the mechanism of force depression

Active shortening against the stretch of series elasticity during force development is ubiquitous in skeletal muscle, yet it is our understanding that this study is the first to directly quantify shortening-induced force depression during a maximal fixed-end contraction (Raiteri & Hahn, 2019; Raiteri et al., 2024). We characterised the effect of added compliance on the depression of maximum isometric force, finding that when fibre shortening against the stretch of a spring was increased by 0.1 *L*_0_, the maximum force that could be generated decreased by 0.07 *P*_0_. When force depression was measured conventionally, using the fitted curves to compare force at a matched length, *L*_0_, we found force depression to be 0.08 *P*_0_ for a 0.1 *L*_0_ increase in fibre shortening. This extent of force depression generally agrees with that measured previously when a relatively slow shortening ramp was imposed upon a frog muscle fibre. Edman and colleagues (1993) imposed shortening equal to approximately 0.24 *L*_0_ and observed force depression at a length close to optimum to be 16%. For this magnitude of shortening, our regression equation predicts force depression at 20%.

We have demonstrated that the force depression resulting from active shortening against series elasticity, when assessed at a similar length, depends strongly on the mechanical work performed during shortening, as is the case for isokinetic or isotonic shortening (Corr and Herzog, 2005; Granzier and Pollack, 1989; Herzog et al., 2000; Van Noten and Van Leemputte, 2011). Recently, the strength of work as a determinant of force depression was questioned (Raiteri et al., 2024), partly because regression analyses from prior studies had included correlated observations. Since the relationship between work and force depression observed for the current study was derived from independent observations, it seems unlikely that the predictive power of work as a determinant of force depression is an artefact.

Conventionally, force depression is characterised from ramp shortening imposed during the plateau of tetanic force. Shortening is initiated with high activation and high force and causes force to fall. In contrast, shortening against series compliance begins at the onset of contraction, when activation and force are low, and continues as force rises. Because the work dependence of force depression at a given final length has been tested rather thoroughly, we can draw similarities between the current experiment and the experiment of Herzog and colleagues (2000) that involved deactivating the muscle to different extents during the period of imposed shortening. The duration of deactivation imposed was long enough that force approached or reached zero, in which case, active shortening then coincided with rising activation and force development. The force depression resulting from these contractions, when plotted against work, were consistent with data points obtained from several other experiments without deactivation. These additional experiments were performed on the same muscle, manipulating either the amplitude or speed of shortening or muscle recruitment to vary work during shortening. The consistency of the work-force depression relationship suggests that activation and force at the onset of shortening do not independently modulate force depression.

A popular explanation for force depression is that the compliance of the actin filament (Huxley et al., 1994; Kojima et al., 1994), and its deformation during contraction, leads to an inhibition of cross-bridge binding in the region of myofilament overlap formed during shortening (Mariéchal & Plaghki, 1979; Herzog et al., 1998). Since force depression is strongly related to work, this mechanism proposes that the size of the newly formed region and the stress imposed as this new region is formed ultimately determine the extent of force depression (Herzog et al., 2000). More recently, this theory has been expanded to include impaired cross-bridge attachment in the old zone of myofilament overlap, as well as a smaller contribution from reduced force per cross bridge (Joumaa et al., 2012). Joumaa and colleagues (2012) speculated that high stresses lead to an inhibition of cross-binding while low stresses cause a transformation to weakly bound states. The length dependence of this phenomenon is thought to reflect length-dependent compliance exhibited by the free actin segment (Higuchi et al., 1995). Van Noten and Van Leemputte (2011) proposed that force depression normalised to work, which they showed to increase exponentially with length, represented actin compliance.

By increasing series compliance, we increased the initial fibre length required for the active length to fall at the optimum length for force generation. Thus, force depression in the current study increased not only in proportion to shortening and work but also initial fibre length and the region of filament overlap newly formed during shortening. An additional consequence of a longer initial fibre length is that force, and presumably actin stress, is greater for any given amount of overlap during shortening. As such, these findings are seemingly consistent with the previously proposed stress-induced cross-bridge inhibition mechanism (Rassier & Herzog, 2004). Because active shortening increased in proportion to force development, the observed force depression might reflect both cross-bridge inhibition and reduced force per cross-bridge according to the proposed stress dependence of cross-bridge impairment (Joumaa et al., 2012).

Although our findings are consistent with stress-induced cross-bridge inhibition, the mechanism underpinning shortening-induced force depression has received much debate (Herzog, 1998; Rassier & Herzog, 2004; Holt & Williams, 2018; Nishikawa et al., 2018). Titin, a molecular spring responsible for the majority of passive tension in muscle fibres (Prado et al., 2005; Ottenheijm et al., 2009), has also been implicated (Rode et al., 2009; Nishikawa et al., 2012; Jeong & Nishikawa, 2023). Titin’s inherent stiffness increases upon activation through the binding of calcium (Labeit et al., 2003; DuVall et al., 2013). Further increases in stiffness may result from Ca^2+^ mediated titin-actin binding and actin-myosin interaction (Labeit et al., 2003; Joumaa et al., 2008; Cornachione & Rassier, 2011) (Leonard & Herzog, 2010; Powers et al., 2014). According to the winding filament theory (Nishikawa et al., 2012), force depression is due to a reduction in titin-based force after active shortening, which acts to slacken titin and counteract the stiffening effects of activation and force generation. We measured a maximum amount of force depression of 0.25 *P*_0_ for contractions ending at, or slightly below, optimum length. Optimum sarcomere length in frog fibres is 2.0-2.2 μm (Gordon et al., 1966; Edman, 1966). Passive tension, however, only begins developing once frog fibres are stretched beyond a length of approximately 2.7 um (ter Keurs et al., 1978; Edman, 1979; Altringham & Bottinelli, 1985). Therefore, it seems that a substantial reduction in titin’s slack length in response to activation is a minimum requirement for any mechanism relating force depression to a reduction in titin-based force.

Activation has a dramatic effect on the apparent stiffness of titin (Leonard & Herzog, 2010; Powers et al., 2014), however, titin’s slack length may not be easily modified. Calcium alone has virtually no effect on fibre slack length (Labeit et al., 2003; Joumaa et al., 2008; Leonard & Herzog, 2010; Cornachione et al., 2016) and exerts only a modest effect on fibre stiffness. It also appears that when myofibrils are activated at optimum length (without inhibition of myosin-actin interaction) and very slowly stretched (< 0.05 *L*_0·_s^-1^), force does not immediately increase (Leonard & Herzog, 2010). A significant increase in force was only evident once myofibrils were actively stretched to a length corresponding to the onset of passive force in resting myofibrils. This observation seems to be consistent with the finding that passive force enhancement — the elevated passive force exhibited by resting muscle after active stretch — is absent on the ascending limb, at optimum length, or at lengths that don’t correspond to passive force in resting muscle (Herzog & Leonard, 2002, 2005; Peterson et al., 2004; Rassier et al., 2005; Hisey et al., 2009). Passive force enhancement is understood to be the passive component of the enhanced force exhibited by actively stretched muscle (i.e., residual force enhancement) and is attributed to increased titin stiffness (Herzog, 2019; Herzog and Leonard, 2005; Rassier and Herzog, 2004). These findings suggest that titin’s slack length isn’t significantly altered by actin-myosin interaction either.

If titin remains slack in active muscle at optimum length, or bears only a small tension when stretched, doubt is raised as to whether titin contributes to the deficit in isometric force at optimum length, let alone lengths on the ascending limb. Our reluctance to implicate titin in force depression seems reasonable since stretches of varying speed and amplitude imposed immediately before active shortening do not influence force depression (Herzog & Leonard, 2000; Lee et al., 2001). Activating titin at unique lengths and stretching titin by different extents would be expected to give unique titin properties at the onset of shortening (Leonard & Herzog, 2010; Herzog, 2014) and presumably afterwards.

## CONCLUSION

We demonstrate that active shortening against series elasticity during a fixed-end contraction induces force depression. The magnitude of force depression can be substantial, increases in proportion to shortening amplitude and mechanical work, and is dependent on length. The interaction of the length and work dependencies of force depression with the length dependence of myofilament overlap causes tetanic optimum length to be reduced when series compliance is added. We propose that significant series compliance increases the activation dependence of optimum length by reducing tetanic optimum length with respect to the length corresponding to optimum filament overlap. Our results indicate that the shift in optimum length mediated by series compliance and force depression is too modest to account for the pronounced activation dependence exhibited by the bullfrog plantaris muscle, and that additional mechanisms should be sought to explain the apparent force dependence of optimum length found for this muscle. Muscle-specific work and length dependencies of force depression should also be considered. The force depression that arises from shortening against series elasticity during force development is consistent with a stress-induced inhibition of cross-bridge binding.

## DATA VAILABILITY STATEMENT

All data for individual muscles supporting major results are included in manuscript figures. Raw recordings of muscle force and fascicle length will be made available upon reasonable request.

## AUTHOR CONTRIBUTIONS

Experiments were performed in the laboratory of N.C.H. at the University of California, Riverside. D.L.M. and N.C.H contributed to conceptualisation, methodology, and interpretation of data; D.L.M. performed experiments, developed software, completed the formal analysis and visualisation of data, and drafted the manuscript; D.L.M. and N.C.H. contributed to the revision and editing of the manuscript. All authors approved the final version of the manuscript and agree to be accountable for all aspects of the work. All persons designated as authors qualify for authorship, and all those who qualify for authorship are listed.

